# Geographic variation in larval cold tolerance and exposure across the invasion front of a widely established forest insect

**DOI:** 10.1101/2021.12.08.471760

**Authors:** Petra Hafker, Lily M. Thompson, Jonathan A. Walter, Dylan Parry, Kristine L. Grayson

## Abstract

As the global climate changes, high and low temperature extremes can drive changes in species distributions. Across the range of a species, thermal tolerance can experience plasticity and may undergo selection, shaping resilience to temperature stress. In this study, we measured variation in the lower thermal tolerance of early instar larvae of an invasive forest insect, *Lymantria dispar dispar* L. (Lepidoptera: Erebidae), using populations sourced from the climatically diverse invasion of the Eastern United States. In two chill coma recovery experiments, we recorded recovery time following a period of exposure to a non-lethal cold temperature. A third experiment quantified growth responses after chill coma recovery to evaluate sublethal effects. Our results indicate that cold tolerance is linked to regional climate, with individuals from cold climate populations recovering faster from chill coma. While this geographic gradient is seen in many species, detecting this pattern is notable for an introduced species founded from a single point-source introduction. We demonstrate that the cold temperatures used in our experiments occur in nature from cold snaps after spring hatching, but negative impacts to growth and survival appear low. We expect that population differences in cold temperature performance manifest more from differences in temperature-dependent growth than acute exposure. Evaluating intraspecific variation in cold tolerance increases our understanding of the role of climatic gradients on the physiology of an invasive species, and contributes to tools for predicting further expansion.

**Summary Statement:** Chill coma recovery experiments demonstrate geographic variation in cold temperature tolerance for a widespread invasive species, and the risk of early season exposure differs across the invasion front.

## Introduction

Global climate change is driving complex and geographically variable alterations to temperature and microclimate, which in turn can impact population persistence and geographic ranges of organisms. These consequences occur through the impacts of climate change on phenology, growing season length, or the frequency and severity of thermal extremes (Yang et al., 2021). Insects, as short-lived ectotherms, are excellent systems for studying these responses as their physiological performance is sensitive to temperature and their small size allows for experimental studies that might not otherwise be feasible in larger organisms. While research on organismal impacts from climate change often focuses on increasing temperatures, the effects of amplified variability in winter and spring temperatures and the changing frequency of thermal extremes are equally important in the responses of organisms (Cohen et al., 2014; Vavrus et al., 2006). For example, weakening of jet stream air flows or a stratospheric polar vortex can result in unusually extreme cold events when arctic air reaches mid-latitudes (Kretschmer et al., 2018), potentially causing high mortality in overwintering or recently hatched insects (MacQuarrie et al., 2019). Reduced snowfall or earlier snowmelt can reduce insulation and decrease microclimate temperatures for soil-dwelling species or life stages (Lawrence & Slater, 2010). Shifts in winter conditions and resilience to cold are key drivers of the impacts to insects from climate change (Marshall et al., 2020; Roland & Matter, 2016; Williams et al., 2015).

Cold stress can cause mortality, as well as sublethal effects on growth and development, for both overwintering insects and larvae at risk of cold snaps after emergence. Many studies have found differences among populations or between closely related species in cold tolerance traits, suggesting the potential for evolutionary responses or plasticity to cold temperatures (Ayrinhac et al., 2004; MacLean et al., 2019). The potential for populations to respond to selective pressure from cold stress is particularly important as species’ ranges can shift, expand, or contract under climate change (Lancaster, 2016; Pureswaran et al., 2018). In these scenarios, insects may experience novel exposure to cold extremes. Invasive species provide a useful surrogate for understanding the evolutionary potential of thermal traits under novel environments in the face of widely changing climate conditions and shifts in species’ ranges (Jactel et al., 2019; Kindlmann et al., 2001). Although many invasive species are generalists that can thrive in a wide range of environments, physiological limits to climate tolerance still ultimately shape their potential range (Huey & Hertz, 1984; Sinclair et al., 2012). Often originating from a single or small number of localized introductions, evolutionary responses during range expansion of invasive species can shed light on the role of plasticity and evolutionary adaptation to extreme temperatures.

Several metrics have been used to compare the ability of insects to withstand low temperatures, including cold hardening, chill coma recovery time, and lower critical thermal limits (Sinclair et al., 2015; Sinclair & Roberts, 2005; Zachariassen, 1985). Chill coma recovery (CCR) is a measure of the time it takes for an individual to regain motor function after being held in low temperature paralysis and returned to an ambient temperature (MacMillan & Sinclair, 2011). This metric has been widely used as a measure of cold tolerance and used in studies examining adaptation to climatic gradients (e.g., Castañeda et al., 2005). Many studies have examined interspecific variation in thermal minimum traits, focusing on macroevolutionary comparisons using insect taxa with species in both tropical and temperate zones to examine the phylogeographic history of cold tolerance traits (David et al., 2003; Gibert & Huey, 2001; Hoffmann et al., 2002). Examining microevolutionary changes that result in intraspecific variation can provide a window into the potential for adaptive responses of a species to climate change and the development of geographic gradients in thermal traits. Variation in cold tolerance traits, especially related to CCR, has been most comprehensively studied in and among *Drosophila* species (e.g., David et al., 1998; Gerken et al., 2016; Gibert et al., 2001; Sisodia & Singh, 2010). In this study, we use the invasion of a well-known forest insect, *Lymantria dispar dispar* L. (Lepidoptera: Erebidae; the previous common name, European gypsy moth, is currently under revision) to test for geographic variation in cold tolerance across its invasive range in Eastern North America.

Since being introduced in North America over 150 years ago, *L. d. dispar* has spread from its introduction point near Medford, Massachusetts, USA to inhabit diverse climatic conditions encompassing cold extremes in Minnesota and Eastern Canada to sub-tropical climates at the current southern extreme in North Carolina (Figure 1; Grayson & Johnson, 2018). Genetic evidence points to a single primary originating population in Massachusetts, which was introduced in 1869 (Wu et al., 2015). Continuous range expansion of *L. d. dispar* over a century allows us to ask questions about the potential for plasticity and evolutionary responses to create intraspecific variation in performance traits among populations, and may inform more broadly, our understanding of trait changes in response to novel climatic environments. Furthermore, invasive species can provide a real-time proxy for the types of temperature shifts facing many other species under anthropogenic climate change.

**Figure 1.**
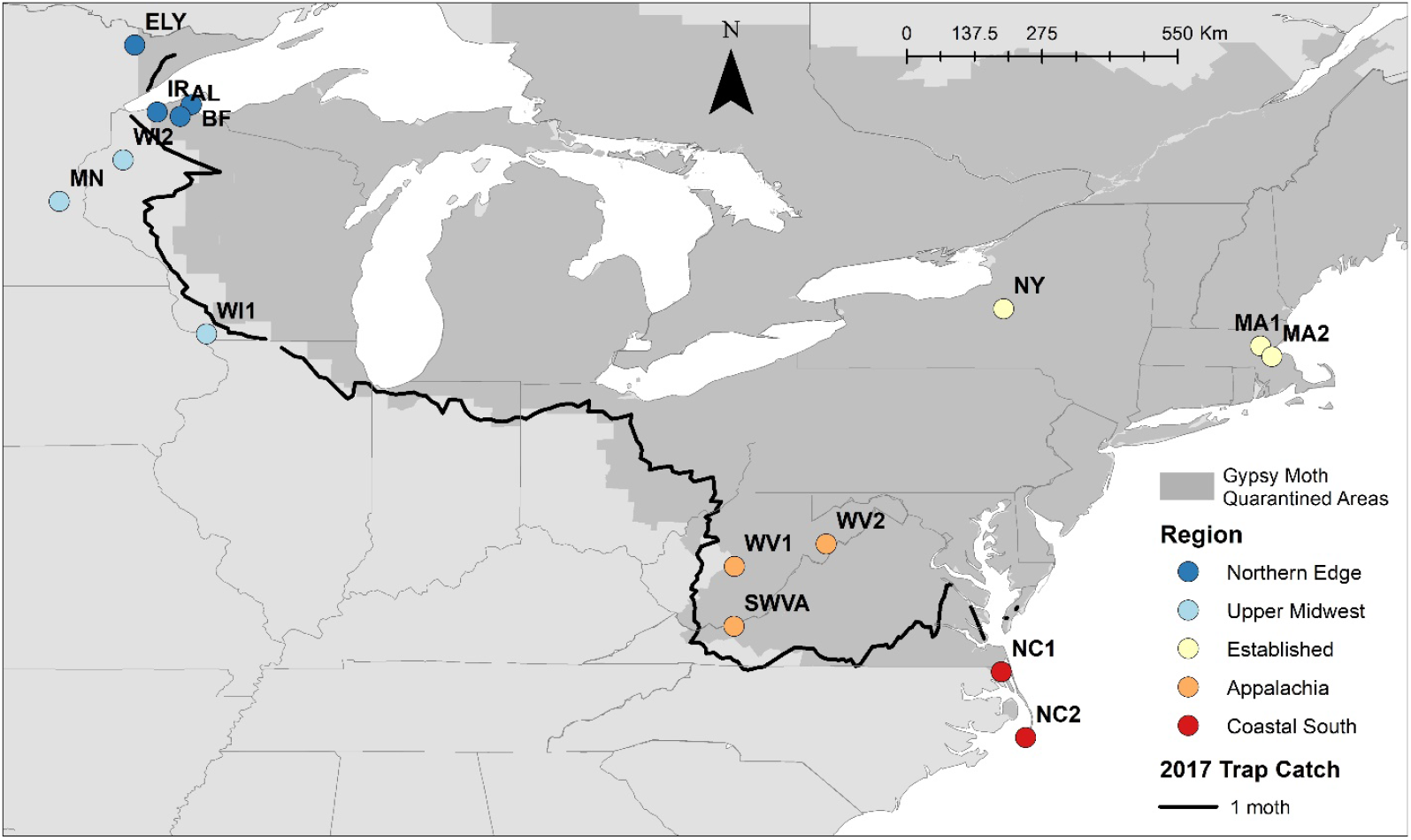
Map of *Lymantria dispar dispar* populations used in this study relative to the invasion front and established range in North America. The established range (darker grey shading) is indicated by counties designated under USDA *L. d. dispar* quarantine regulations. The invasion front shown is the 1-moth trap catch line from the Slow-the-Spread program (Sharov et al., 2002) in 2017. Source populations are colored by the regions defined in this study.

In this study, we measured CCR in 15 populations of *L. d. dispar* sourced from the extremes of the United States invasive front as well as from long established areas near the introduction site. We tested whether caterpillar larvae from northern *L. d. dispar* populations are more tolerant of cold temperature stress than those from southern portions of the invasive range by comparing larval CCR among populations for two early instar stages. In addition, we quantified sublethal effects of cold exposure in a post-chill coma growth experiment and determined the magnitude of cold exposure that each population of larvae experience in their source environment in the spring after hatching. Given previous research suggesting local adaptation has occurred in thermal performance traits and heat tolerance across the *L. d. dispar* invasive range (Thompson et al., 2017, 2021), we predicted that individuals from northern populations would recover from a chill coma more quickly than those from southern populations. Similarly, we predicted a lag effect on larval growth of *L. d. dispar* after recovery from chill coma, and that this effect would be greater in populations from southern sources than in the north. Investigating intraspecific variation in thermal tolerance will increase our understanding of how these traits can change during the course of an invasion, and whether climate change may alter the success and future spread of this economically and ecologically harmful *s*pecies in North America.

## Materials and Methods

### Study system

*Lymantria dispar dispar* L. (Lepidoptera: Erebidae) is a generalist-defoliating forest pest that was introduced to North America in 1869. Populations of this species experience cyclical outbreaks that can cause extensive ecological and economic damage to hardwood forests and disrupt ecosystem processes (Clark et al., 2010; Elkinton & Liebhold, 1990). The life cycle of *L. d. dispar* includes obligate diapause and overwintering as egg masses before hatching in the spring, typically at or near bud break of deciduous trees. The species is a broadly generalist foliage feeder and undergoes five to six larval instars before pupation (Liebhold, 1995). Adult moths are sexually dimorphic, short-lived, and do not feed. Females of this subspecies are flightless and use pheromones to attract flying males for reproduction. Spread of *L. d. dispar* can be primarily attributed to larval ballooning and human-mediated movement of egg masses attached to objects, vehicles, and firewood (Doane & McManus, 1981).

We used egg masses collected in the field from 15 populations representing five geographic regions in the Eastern United States, including the established invaded range and zones of the invasion front (Figure 1). Egg masses were collected from the field in 2017 or earlier (see Thompson et al., 2021) and successive generations were reared outdoors under identical conditions on oak foliage near Syracuse, NY until shipped to Richmond, VA as egg masses for these experiments (USDA APHIS permits P526P-17-03681(KLG) and P526P-16-04388 (DP)). For the invasion front regions, four populations were sourced from the extreme **northern edge** (AL, BF, IR, ELY) and three populations were from the **upper Midwest** (WI1, WI2, and MN). In the southern portion of the invasion front, three populations came from **Appalachia** (WV1, WV2, and SWVA) and two populations from the **coastal south** in North Carolina (NC1 and NC2). From the **established** invaded range, three populations were sourced from the early invaded range (MA1, MA2, and NY), including two in close geographic proximity to the species’ introduction point in Medford, MA (MA1 and MA2). Multiple populations from the five delineated regions were used to account for possible maternal effects from variation between populations in host foliage quality (Erelli & Elkinton, 2000; Rossiter et al., 1993) and to increase the generality of the results for a given geographic region.

### CCR Experiment 1

We first measured CCR in 2018 using third instar larvae from 14 populations. This initial experiment used 2°C for 20 hours as the cold exposure and scored larval recovery time by direct observation from a single observer. The populations tested and sample sizes are reported in Table 1.

**Table 1.** Population abbreviations, regions, and sample sizes used in the experiments in this study.

| Region | Population | Elevation<br>(m asl) | Sample Size |  |  |
| --- | --- | --- | --- | --- | --- |
|  |  |  | *CCR 1* | *CCR 2* | *Growth* |
| Northern<br>Edge | ELY | 446 |  | 25 | 40 |
|  | BF | 222 | 24 | 32 | 39 |
|  | IR | 269 | 21 | 32 |  |
|  | AL | 297 | 30 | 17 |  |
| Upper<br>Midwest | WI2 | 353 | 22 | 31 | 39 |
|  | MN | 261 | 19 |  |  |
|  | WI1 | 243 | 25 |  |  |
| Established | NY | 146 | 18 | 26 |  |
|  | MA1 | 56 | 23 | 33 |  |
|  | MA2 | 96 | 21 | 26 | 38 |
| Appalachia | WV2 | 525 | 20 |  |  |
|  | WV1 | 614 | 19 | 23 |  |
|  | SWVA | 528 | 20 | 30 | 32 |
| Coastal<br>South | NC1 | 3 | 35 | 45 |  |
|  | NC2 | 1 | 28 | 36 | 36 |

After hatching, larvae were reared in an environmental chamber at 25°C (Model DROS33SD, Powers Scientific Inc., Doylestown, PA, USA) and fed a wheat-germ based artificial diet (USDA APHIS Formulation). Upon reaching third instar, larval mass was recorded to the nearest thousandth of a gram and individuals were placed in 30 ml clear, lidded cups without diet. These cups were then transferred into a dark, 2°C chamber for 20 hours (Model I-22VL, Percival Scientific, Inc., Perry, Iowa, USA). The temperatures in the chamber and in the lab were recorded every 10 minutes with data loggers to ensure that chilling conditions were maintained throughout the 20-hour period (HOBO U23 Pro v2, Onset Computer Corporation, Bourne, MA, USA). Following the cold exposure, approximately five caterpillars were removed from the chamber at a time, according to staggered chill coma start time, and placed on a lab bench at ambient room temperature. A single observer monitored larval movement at eye-level, six inches from the cups. The time in seconds was recorded for each individual larva to regain motor function (e.g., curling of the head or abdomen, twitching movements, movement of legs or prolegs). The time frame began with entry into the room temperature environment and ended with the first indication of movement. Individuals were included in the analysis if movements were visible in real-time to the naked eye (15 individuals were excluded for failing to meet this criteria, no more than four per population).

### CCR Experiment 2

We conducted a second CCR experiment in 2020, this time using second instar larvae with a cold exposure of 1°C for 20 hours to test a colder temperature at a life stage closer to spring hatching. We had 13 populations available for this experiment (see Table 1 for sample sizes). Larval recovery was observed for up to 54 individuals at a time by two observers and video recordings were used to verify the scored times.

After hatching, larvae for this experiment were reared in environmental chambers at 24°C (Model I-22VL, Percival Scientific, Inc., Perry, Iowa, USA) on the same wheat-germ based diet as in CCR Experiment 1. Given that second instar larvae are very small, mass measurements at the thousandth of a gram did not provide enough precision to detect differences between individuals. Instead, we standardized the developmental point of individuals used in this experiment as two days post-molt.

To reach a colder exposure temperature, we used a benchtop chiller with an antifreeze bath set to 1°C (MicroCooler II Model 260010, Boekel Scientific, Feasterville-Trevose, PA, USA). Second instar larvae were placed into microcentrifuge tubes with three or fewer larvae from the same population for 20 hours. An empty vial with a datalogger probe monitored temperature during the trial (EasyLog Data Logger EL-USB-TP-LCD, Lascar Electronics, Erie, PA, USA).

After removal from the chiller, larvae were directly placed in labeled petri dishes for each population and recorded using a webcam (Model C910 Logitech HD Pro Webcam, Logitech, Lausanne, Switzerland). One live observer and one virtual observer were present for each trial due to Covid-19 pandemic restrictions in 2020. Recovery time was scored based on observations of first movement, as defined in CCR Experiment 1 (Video S1). Observed movements were verified from the video recording and recovery time was calculated from the time individuals were removed from the chiller and placed in the observation dish. Individuals that crawled into the lid of the microcentrifuge tube had refuge above the liquid level of the chiller and were excluded (13 individuals were excluded, no more than four per population).

### Post-Chill Coma Growth Experiment

We designed an additional experiment to measure the effects on subsequent growth following a cold exposure resulting in a chill coma. Concurrent with CCR experiment 2 in 2020, second instar larvae from six populations (Table 1) were exposed to a chill temperature or control treatment for 24 hours before being returned to normal diet and rearing temperature. Survival and mass were assessed 10 days post-treatment. Measuring growth over a longer duration was not possible due to Covid-19 pandemic restrictions that limited the number of lab personnel onsite.

Forty larvae were reared from hatch on the same wheat-germ based diet as above in an environmental chamber at 24°C (Model I-22VL, Percival Scientific, Inc., Perry, Iowa, USA). Starting size of larvae in the experiment was standardized by using individuals one day following molt. Ten individuals from each population were randomly assigned one of four treatment groups for 24 hours (Figure 2): 1) remain at 24°C on diet (control treatment), 2) placed in an empty cup at 24°C without diet (a starvation control), 3) chilled at 1°C (cold exposure without food), 4) chilled at -2.5°C (cold exposure without food). Larvae in the chilled treatments were placed in the same vials and chiller as in CCR experiment 2. The starvation control at 24°C allowed us to separate the effects of the cold exposure treatments from the effect of being removed from food. Following the 24-hour treatment period, individuals were returned to individual containers with diet and the 24°C environmental chamber. After ten days, survival was assessed and the mass of each surviving individual was recorded to the nearest thousandth. Individuals that were dead at 10 days post-treatment (16 out of 240) were excluded from the analysis (Table 1).

**Figure 2.**
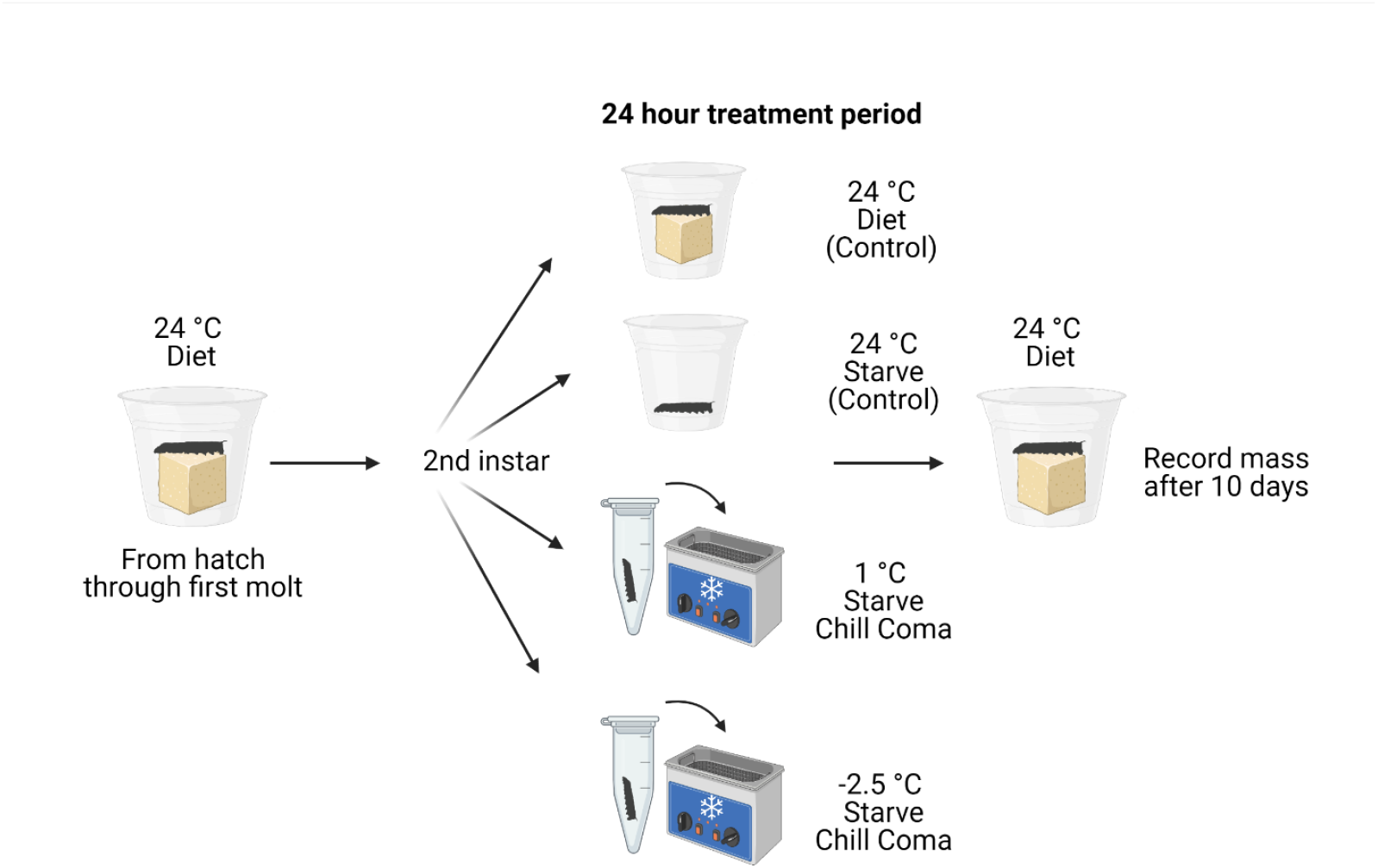
Design of the post-CCR growth experiment. All individuals were reared to the second instar on artificial diet before being assigned to one of four treatment groups for 24 hours (N = 40 individuals each for six populations). Following exposure to the treatments, all individuals were returned to initial rearing conditions. The mass of each surviving individual was recorded after a 10 day post-treatment growth period. Created with BioRender.com.

### Experimental Data Analysis

We examined patterns in CCR across populations relative to the climate of each of the source locations as in Thompson et al. (2021). We examined differences between populations using linear regression with CCR in seconds as the dependent variable and the mean 30-yr climate normal (1981-2010) of the population source location as the independent variable (PRISM Climate Group, Oregon State). With multiple populations represented per region, this approach allowed us to appropriately scale populations based on differences in thermal environments more directly than using a proxy metric such as latitude. For CCR experiment 1, individual mass (g) was included as a covariate while size was standardized in experiment 2 by days post-molt. In the statistical analyses, both climate normal and mass variables were scaled to a mean of zero and standard error of one to account for the difference in magnitude of the values.

In the growth experiment, differences in mass of larvae after exposure to the treatments was analyzed using a two-way ANOVA. The mass (g) of individuals 10 days after treatment exposure was the dependent variable (starting mass was too small to detect differences between individuals at a thousandth of a gram) and the independent variables were the treatment, the population, and their interaction. A post-hoc Tukey Honest Significant Differences test was conducted for the significant independent variables in order to identify the effects that contributed most to differences in mass.

To meet the assumptions of normality, CCR values in CCR experiment 1 were log-transformed. Tansformation of the dependent variable was not necessary for CCR experiment 2 or the growth experiment. Analyses for all experiments were conducted in Program R (version 3.6.2; R Core Team, 2019) and significance was evaluated at *α*= 0.05.

### Site temperatures

Finally, we aimed to contextualize our experiments in terms of cold temperatures experienced by wild populations of *L. d. dispar* during larval development. To do this, we quantified the number of days with minimum temperatures below thresholds from the temperatures we used in our experiments (2°C, -2.5°C). Using the locations of the source populations, we quantified the amount of cold each year from the projected median egg hatch date until the end of adult development.

We extracted a daily temperature time series for 2000 to 2020 for the locations of all 15 source populations using DAYMET data (Thornton et al., 2020) accessed through the ‘daymetr’ R package (Hufkens et al., 2018). Median egg hatch date for each year of the temperature time series was projected using the egg development model of the *L. d. dispar* life stage model (GLS; Gray, 2009). The number of days during larval development (as defined by the median egg hatch date + 90 days) with minimum temperatures below each threshold was totaled for each year and source population location, then summarized by region.

## Results

### CCR Experiment 1

All individuals used in the first CCR experiment survived and regained motor function following chill coma. During the experiment, the environmental chamber had a mean temperature of 2.4°C (±0.007°C SE) and the ambient room temperature ranged from 22.7°C to 26.4°C. Mean population CCR after a 20-hour exposure to 2°C for the third instar larvae ranged from 220 to 1005 seconds. There was a positive relationship between the 30-year climate normal for a source population location and mean CCR (Adj. R^2^ = 0.27, intercept = 5.91, slope for climate = 0.36, F_(2, 322)_ = 61.7, p < 0.001; Figure 2A); populations from warmer climates took longer to recover from chill coma than populations from colder climates.

### CCR Experiment 2

All individuals also survived and regained motor function in CCR experiment 2. Two chillers were used and the mean temperature for each chiller across all trials was 1.30°C and 1.32°C with a standard error of ±0.006°C and ±0.010°C, respectively. The ambient temperature was recorded at the start of the CCR period and ranged from 23.7°C to 25.2°C across all trials. Mean population CCR of second instar larvae ranged from 723 to 1026 seconds following a 20-hour exposure to 1°C. A weak positive relationship was found between the 30-year climate normal and mean CCR (Adj. R^2^ = 0.043, intercept = 733.1, slope = 12.5, F_(1, 354)_ = 13.5, p < 0.001; Figure 2B); again, populations from warmer climates tended to take longer to recover from chill coma.

### Post-Chill Coma Growth Experiment

The temperatures of the 1°C and -2.5°C cold environment chillers were recorded every two minutes yielding mean (±SE) temperatures of 1.22 ± 0.005°C and -2.95 ± 0.013°C. Survival ranged from 80% to 100% for the 40 larvae in each population and treatment combination (Table 1). Occasional morality was seen in all treatments and populations, with slightly lower survival in the Appalachia and Coastal South populations (SWVA and NC2, respectively).

The interaction of population and treatment did not significantly contribute to the observed differences in growth after chill coma (F_(15, 200)_ = 1.4, p = 0.14). Treatment (24°C Diet; 24°C Starve; 1°C Starve; -2.5°C Starve) significantly predicted individual mass 10 days post-treatment (F_(3, 200)_ = 27.83, p < 0.001), with the control treatment (24°C, Diet) resulting in the largest mass and the coldest treatment (−2.5°C, Starve) having the smallest mass (Figure 3). All treatments were significantly different from each other (all p <0.007) based on a pairwise Tukey HSD test except for the 1°C Starve treatment and the 24°C Starve treatment (p = 0.96). Population was also a significant predictor of individual mass 10 days post-treatment (F_(5, 200)_ = 13.06, p < 0.001). Based on a pairwise Tukey HSD test, the WI2 population was significantly different from all other populations (p < 0.001), but the rest of the populations were not significantly different from each other (p > 0.05; Figure 3).

**Figure 3.**
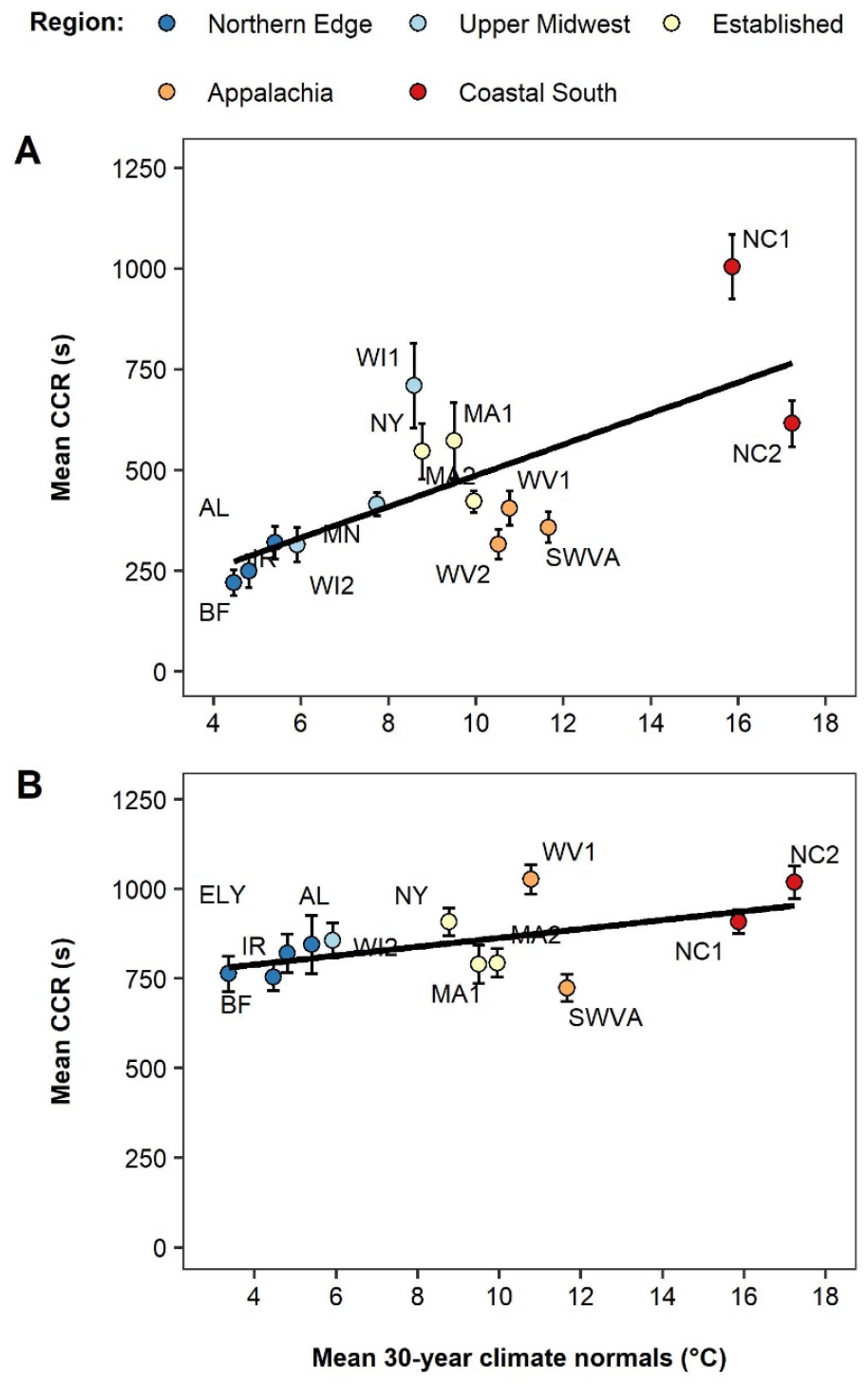
The relationship between population climate normals and CCR time for experiments 1 (A) and 2 (B). Experiment 1 tested 3rd instar caterpillar larvae chilled at 2°C for 20 hours. Experiment 2 tested 2nd instar larvae chilled at 1°C for 20 hours. The specific populations used in each experiment and sample sizes are listed in Table 1.

**Figure 4.**
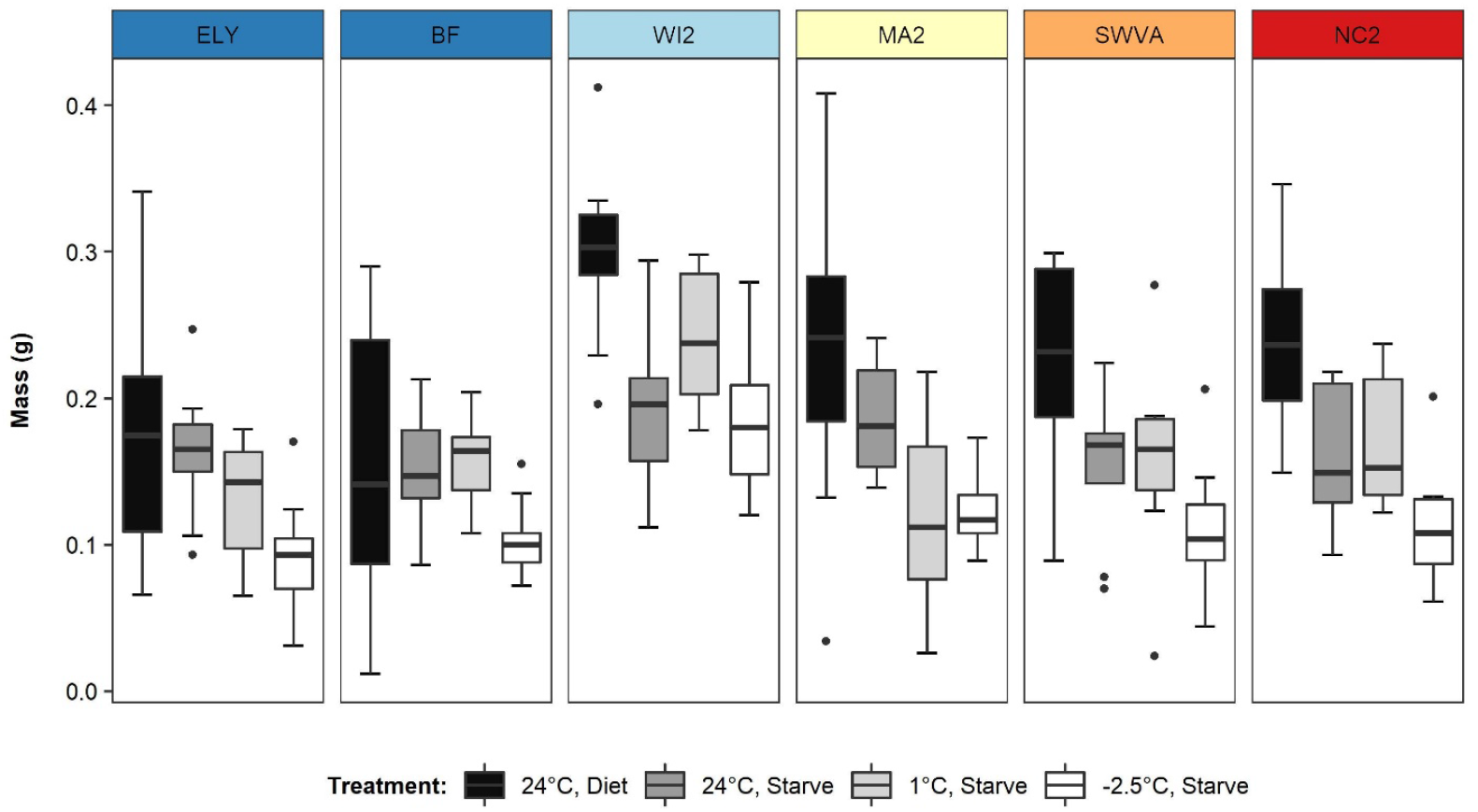
Boxplots of treatment medians per population in the post-chill coma growth experiment. The 24-hour treatments included two controls at 24°C (with artificial diet and starved) and two chilled treatments (at 1°C and at -2.5°C; illustrated in Figure 2). Mass was measured after 10 days of growth post-exposure to the treatments. Box limits indicate the 25th and 75th percentiles of the mass data and bars extend 1.5 times the interquartile range from the 25th and 75th percentiles. Outliers are represented by dots.

### Site Temperatures

The 15 population locations in this study are representative of both long-established and recently expanding areas of the *L. d. dispar* range. Between 2000 and 2020, all populations experienced years with at least one day where the minimum temperature was < 2 °C after predicted larval hatch and during larval development. The southernmost population considered in this study (NC2) was the only location not to experience at least one year with a day where minimum temperature < -2.5 °C during predicted larval development (Figure 5, Table S1). The number of days each year with minimum temperatures below -2.5°C during predicted larval development ranged from 0 to 26 for each site and the number of days below 2°C ranged from 0 to 48 days (Figure 5). Population locations in the Appalachian region experienced the largest median number of cold days during predicted larval development across years (Figure 5, Table S1).

**Figure 5.**
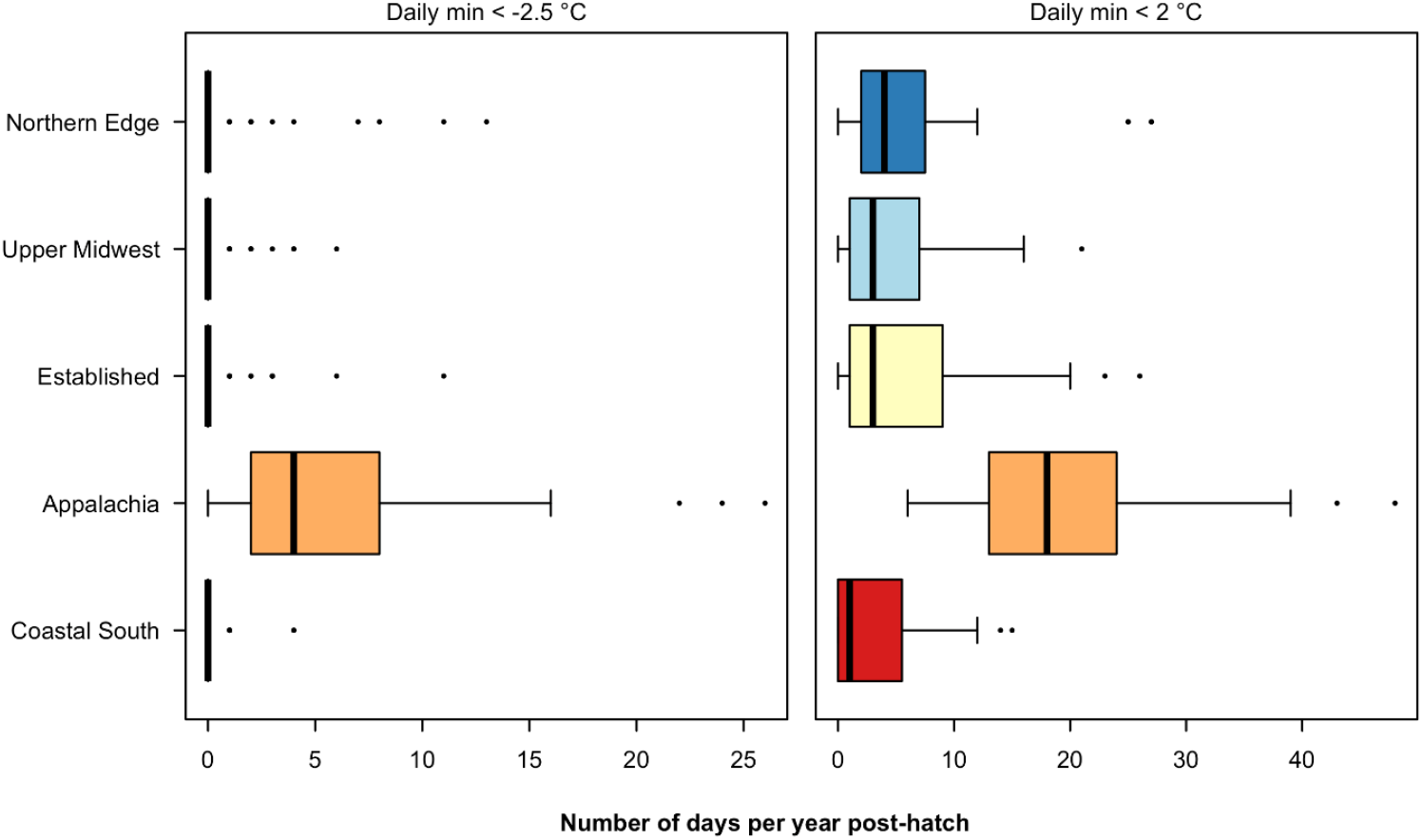
Boxplots of the median number of cold days per year (2000 to 2020) during *Lymantria dispar dispar* larval development by region. Cold days were defined as days with minimum temperatures below -2.5°C and 2°C after the projected spring hatch of larvae each year. There are 2 to 4 sites per region corresponding to locations of source populations, and regions are ordered from lowest mean annual temperature to warmest. Box limits indicate the 25th and 75th percentiles of the mass data and bars extend 1.5 times the interquartile range from the 25th and 75th percentiles. Outliers are represented by dots.

## Discussion

This study tested for geographic variation in cold temperature tolerance of an invasive forest insect using CCR experiments. This well-known metric is used for comparative research on cold tolerance and a wide range of studies have used CCR to examine evolutionary and geographic patterns in temperature performance between species or populations (e.g., Klock & Chown, 2003; MacMillan & Sinclair, 2011). Our study demonstrates that geographic patterns in CCR can also develop across populations of an invasive species with an expanding range. We found that populations of *L. d. dispar* from colder climates have shorter CCR times than those from warmer climates. We quantified the frequency of early season cold exposure experienced by larval *L. d. dispar* in nature and found that spring cold snaps are common for this species after hatching. Exposure to the chill coma-inducing temperatures used in this study did not impact larval survival, but a single cold exposure did have sublethal effects on growth for larvae chilled to -2.5°C for 24 hours. Exposure to 1°C was found to be comparable to larvae removed from food for a similar duration. Under changing climates, resilience to cold temperatures may have important consequences for the development of insect species with early spring hatching phenology.

In our CCR experiments, populations of *L. d. dispar* from warmer climates took longer to recover from exposure to cold temperatures and these geographic differences are consistent with local adaptation of low temperature tolerance traits in colder climates. The potential for intraspecific variation and the genetic basis for CCR response has been demonstrated in several insect species. Similar studies of variation in CCR of wild-sourced populations of *Porcellio laevis* (Castañeda et al., 2005) and *Eldana saccharina* (Kleynhans et al., 2014) also found that populations from warmer locations took longer to recover from chill coma than those from locations with cooler climates. Heritability of cold tolerance has been extensively studied within and among *Drosophila* species (Morgan & Mackay, 2006), where artificial selection has demonstrated evolutionary changes in cold-susceptible traits including CCR response (Gerken et al., 2016). While heritability of cold tolerance has not been examined in *L. d. dispar*, previous work has documented genetic variation between populations (Wu et al., 2015) and spatial genetic structure in development traits (Friedline et al., 2019). Geographic differences in development and growth in response to high temperatures also suggest that local adaptation in thermal traits is occurring in populations across the invasive range (Faske et al., 2019; Thompson et al., 2017).

The geographic pattern in CCR was observed in two experiments that tested different instars and different chill-coma inducing temperatures. New equipment available in 2020 allowed testing a colder exposure temperature and we used the opportunity to test both an earlier life stage and more extreme temperature in our second experiment. We found generally shorter recovery times (i.e., higher cold tolerance) for third instar individuals (CCR Experiment 1) than second instar individuals (CCR Experiment 2), but the differences in exposure temperature between the two experiments complicates this comparison. Stage-specific variation has been found in *L. d. dispar* for tolerance to extreme high temperatures (Banahene et al., 2018).

Additional experiments testing the effects of life stage and low temperature exposure in a factorial design would improve our understanding of stage-specific cold tolerance in *L. d. dispar*. Understanding the cold performance of *L. d. dispar* and the potential for local adaptation in tolerance limits is particularly relevant as this invasive species has begun expanding into northern regions with low predicted climate suitability (Gray, 2004). Undoubtedly, winter severity provides the most extreme levels of cold exposure to egg masses and insulation below the snow line has been shown to be a critical factor for survival of overwintering egg masses (Streifel et al., 2019). For larvae, the greatest risks of acute cold exposure come from early spring cold snaps. Using the detailed life stage model available for *L. d. dispar*, we were able to quantify the yearly number of days below 2°C and -2.5°C after the projected spring hatch and found these temperatures to be common in northern and high elevation locations, and even occasionally occurring in southern populations. Years with cold temperatures after hatching are particularly common for our population locations in Appalachia, the region with the most topographical variation. Here, emergence at high elevation locations was predicted to occur more frequently prior to the end of spring freezing events. The exposure temperatures included in our CCR and growth experiments, while being partly determined by the functional limits of the available equipment, are also biologically relevant to cold temperature stress experienced in nature for the study species.

The cold temperatures used in our experiments caused only minor mortality, which suggests that common spring cold snaps of this magnitude would not result in an appreciable reduction in population size or local extinction. Our growth experiment examined whether acute cold exposure in early instar larvae had persistent effects on growth and development, or whether the exposure was no different than being removed from food for an equivalent time period. The lesser level of cold (1°C) resulted in growth levels generally similar to starvation at 24°C, but more extreme cold (−2.5°C) led to moderately reduced growth during the following 10 days. This effect was seen even after just one instance of cold exposure, and our climate data from the source locations indicates that cold exposure can occur repeatedly in early spring for some years in colder locations. Contrary to expectations, this response did not differ between populations in the growth experiment. Based on our past work, population variation in *L. d. dispar* thermal performance during the larval period seems to occur through sublethal effects on temperature-dependent growth rather than acute exposure. Preliminary work indicates that *L. d. dispar* larvae can survive short cold extremes up to -8°C (*KLG unpublished data*), well below typical temperatures in a spring cold snap. Similarly, research on *L. d. dispar* heat stress found tolerance limits well above high temperatures experienced in nature during larval development (Banahene et al., 2018). Thus, sublethal effects of non-optimal, but nonlethal, temperatures can play an important role in the population performance and perhaps, ultimately the realized range of this species. Future work that incorporates longer development times and direct measurement of fitness metrics would improve our knowledge on the constraints and limits of cold temperature exposure on this species.

Our work adds to the growing evidence that as an introduced species spreads across a landscape, selection and plasticity in response to novel climates can result in climate-related variation in physiological traits. Exposure to extreme temperatures can be highly variable across heterogeneous landscapes and growing seasons among years. Complex interactions between climate change and phenology can lead to dramatic shifts in local conditions and mismatches that can influence the exposure of insects to spring temperature extremes after hatching or emergence (Forrest, 2016; Pureswaran et al., 2018; Yang et al., 2021). Invasive species with distributions extending into novel climate regions provide an important example of how temperature constraints can shape thermal performance, and ultimately impact population persistence and range limits.

## Acknowledgements

We thank Trevor Faske (University of Nevada, Reno), Chelsea Jahant-Miller (SUNY-ESF), Ksenia Onufrieva (Virginia Tech), Chris Foelker (WI DATP), Kimberly Theilen-Cremers and Natasha Northrop (MN DA), Chris Elder (NC Dept. of Ag and Consumer Services), Jeff Johnson, Scott Hoffman, and CJ Campbell (WV Department of Agriculture), and Valerie Huelsman (Prince William County VA Public Works) for helping us acquire the egg masses that were used in this study. Thanks to Hannah Coovert for helpful comments during the writing process. This study was conducted under USDA APHIS permit numbers P526P-17-03681 and P526P-20-02026 (KLG).

## Data availability

Data and code accompanying this study will be publicly archived on Dryad upon manuscript acceptance.

## Supplementary Information

**Table S1.**
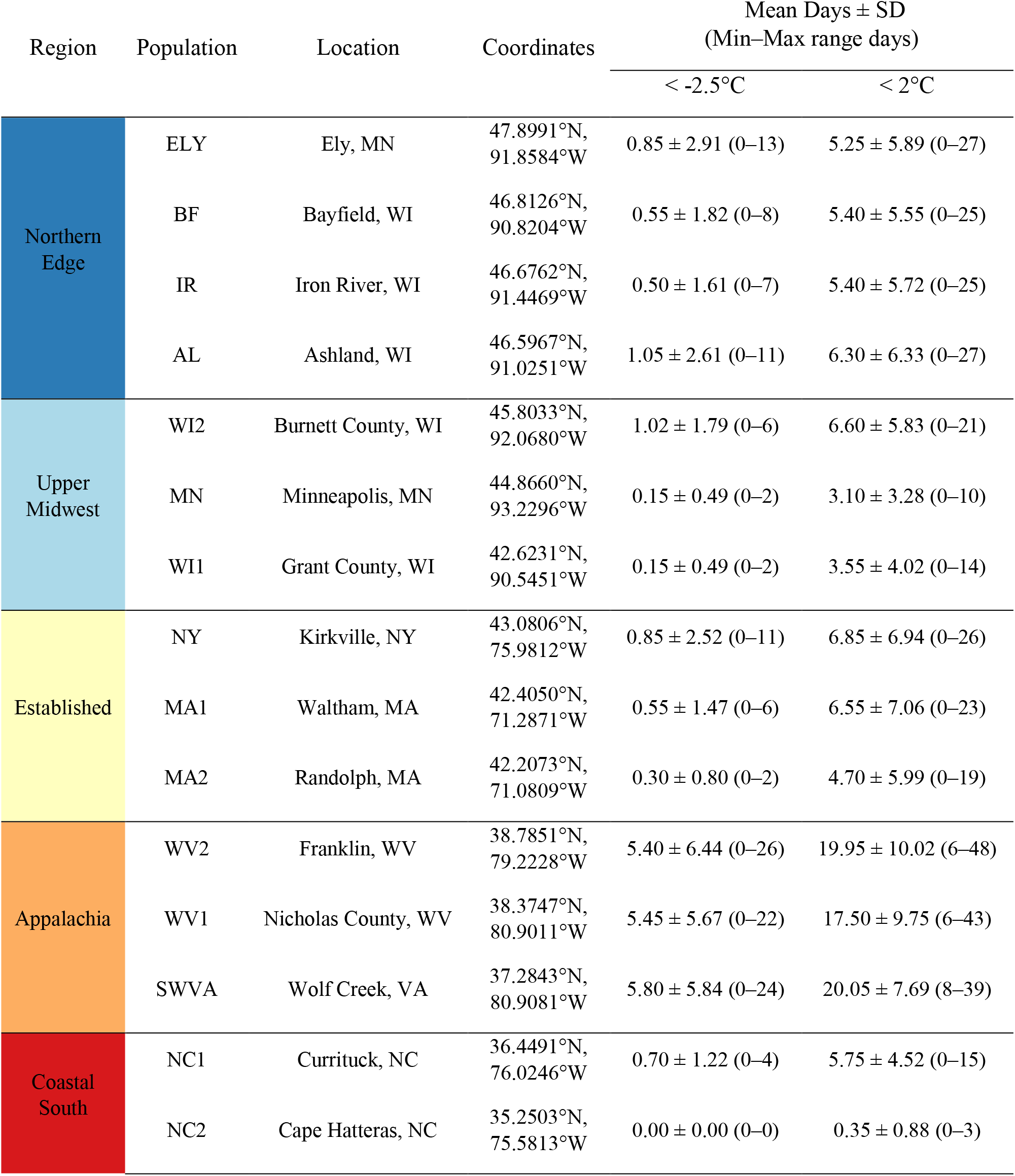
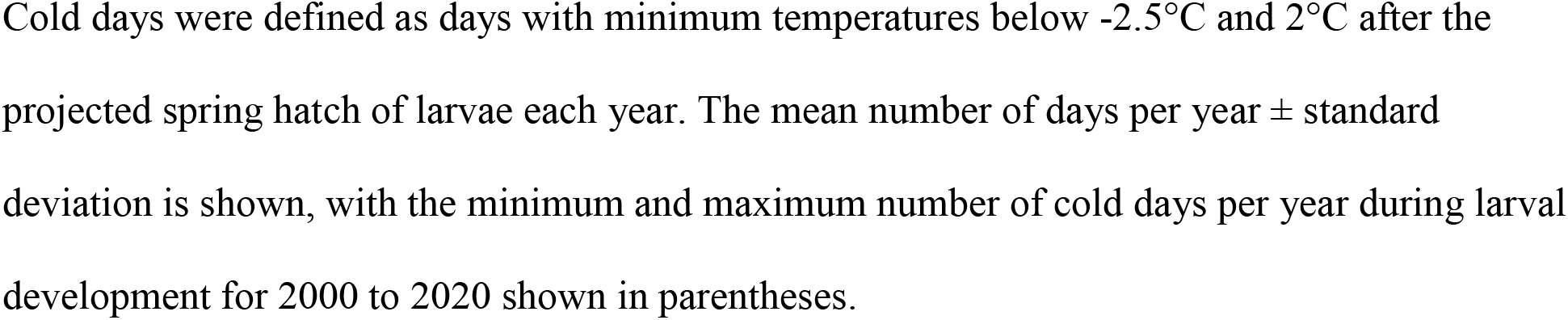
Population location information and descriptive calculations for number of cold days per year (2000 to 2020) for each location during *Lymantria dispar dispar* larval development.

**Video 1**

**Example of recovery during 2020 chill coma recovery (CCR) experiment**. At timestamp 00:12, the abdomen of the second instar individual curls signifying recovery from chill coma recovery. Larvae that have not recovered are positioned ventral side up. Movement for the 2020 CCR experiment is noted as a twitch, or the curling of the head or abdomen. https://youtu.be/1nuACdMZoUE

## References

Ayrinhac, A., Debat, V., Gibert, P., Kister, A.-G., Legout, H., Moreteau, B., Vergilino, R., & David, J. R. (2004). Cold adaptation in geographical populations of Drosophila melanogaster: Phenotypic plasticity is more important than genetic variability. Functional Ecology, 18(5), 700–706. https://doi.org/10.1111/j.0269-8463.2004.00904.x

Banahene, N., Salem, S. K., Faske, T. M., Byrne, H. M., Glackin, M., Agosta, S. J., Eckert, A. J., Grayson, K. L., & Thompson, L. M. (2018). Thermal sensitivity of gypsy moth (Lepidoptera: Erebidae) during larval and pupal development. Environmental Entomology. https://doi.org/10.1093/ee/nvy149

Castañeda, L. E., Lardies, M. A., & Bozinovic, F. (2005). Interpopulational variation in recovery time from chill coma along a geographic gradient: A study in the common woodlouse, Porcellio laevis. Journal of Insect Physiology, 51(12), 1346–1351. https://doi.org/10.1016/j.jinsphys.2005.08.005

Clark, K. L., Skowronski, N., & Hom, J. (2010). Invasive insects impact forest carbon dynamics. Global Change Biology, 16(1), 88–101. https://doi.org/10.1111/j.1365-2486.2009.01983.x

Cohen, J., Screen, J. A., Furtado, J. C., Barlow, M., Whittleston, D., Coumou, D., Francis, J., Dethloff, K., Entekhabi, D., Overland, J., & Jones, J. (2014). Recent Arctic amplification and extreme mid-latitude weather. Nature Geoscience, 7(9), 627–637. https://doi.org/10.1038/ngeo2234

David, R. J., Gibert, P., Moreteau, B., Gilchrist, G. W., & Huey, R. B. (2003). The fly that came in from the cold: Geographic variation of recovery time from low-temperature exposure in Drosophila subobscura. Functional Ecology, 17(4), 425–430. https://doi.org/10.1046/j.1365-2435.2003.00750.x

David, R. J., Gibert, P., Pla, E., Petavy, G., Karan, D., & Moreteau, B. (1998). Cold stress tolerance in Drosophila: Analysis of chill coma recovery in D. melanogaster. Journal of Thermal Biology, 23(5), 291–299. https://doi.org/10.1016/S0306-4565(98)00020-5

Doane, C. C., & McManus, M. L. (1981). The Gypsy Moth: Research Toward Integrated Pest Management. U.S. Department of Agriculture.

Elkinton, J. S., & Liebhold, A. M. (1990). Population dynamics of gypsy moth in North America. Annual Review of Entomology, 35(1), 571–596. https://doi.org/10.1146/annurev.en.35.010190.003035

Erelli, M. C., & Elkinton, J. S. (2000). Maternal effects on gypsy moth (Lepidoptera: Lymantriidae) population dynamics: A field experiment. Environmental Entomology, 29(3), 476–488. https://doi.org/10.1603/0046-225X-29.3.476

Faske, T. M., Thompson, L. M., Banahene, N., Levorse, A., Quiroga Herrera, M., Sherman, K., Timko, S. E., Yang, B., Gray, D. R., Parry, D., Tobin, P. C., Eckert, A. J., Johnson, D. M., & Grayson, K. L. (2019). Can gypsy moth stand the heat? A reciprocal transplant experiment with an invasive forest pest across its southern range margin. Biological Invasions, 21(4), 1365–1378. https://doi.org/10.1007/s10530-018-1907-9

Forrest, J. R. (2016). Complex responses of insect phenology to climate change. Current Opinion in Insect Science, 17, 49–54. https://doi.org/10.1016/j.cois.2016.07.002

Friedline, C. J., Faske, T. M., Lind, B. M., Hobson, E. M., Parry, D., Dyer, R. J., Johnson, D. M., Thompson, L. M., Grayson, K. L., & Eckert, A. J. (2019). Evolutionary genomics of gypsy moth populations sampled along a latitudinal gradient. Molecular Ecology, 28(9), 2206–2223. https://doi.org/10.1111/mec.15069

Gerken, A. R., Mackay, T. F. C., & Morgan, T. J. (2016). Artificial selection on chill-coma recovery time in Drosophila melanogaster: Direct and correlated responses to selection. Journal of Thermal Biology, 59, 77–85. https://doi.org/10.1016/j.jtherbio.2016.04.004

Gibert, P., & Huey, R. B. (2001). Chill-coma temperature in Drosophila: Effects of developmental temperature, latitude, and phylogeny. Physiological and Biochemical Zoology, 74(3), 429–434. https://doi.org/10.1086/320429

Gibert, P., Moreteau, B., Pétavy, G., Karan, D., & David, J. R. (2001). Chill-coma tolerance, a major climatic adaptation among Drosophila species. Evolution, 55(5), 1063–1068. https://doi.org/10.1111/j.0014-3820.2001.tb00623.x

Gray, D. R. (2009). Age-dependent postdiapause development in the gypsy moth (Lepidoptera: Lymantriidae) life stage model. Environmental Entomology, 38(1), 18–25. https://doi.org/10.1603/022.038.0104

Grayson, K. L., & Johnson, D. M. (2018). Novel insights on population and range edge dynamics using an unparalleled spatiotemporal record of species invasion. Journal of Animal Ecology, 87(3), 581–593. https://doi.org/10.1111/1365-2656.12755

Hoffmann, A. A., Anderson, A., & Hallas, R. (2002). Opposing clines for high and low temperature resistance in Drosophila melanogaster. Ecology Letters, 5(5), 614–618. https://doi.org/10.1046/j.1461-0248.2002.00367.x

Huey, R. B., & Hertz, P. E. (1984). Is a jack-of-all-temperatures a master of none? Evolution, 38(2), 441–444. https://doi.org/10.2307/2408502

Hufkens, K., Basler, D., Milliman, T., Melaas, E. K., & Richardson, A. D. (2018). An integrated phenology modelling framework in r. Methods in Ecology and Evolution, 9(5), 1276–1285. https://doi.org/10.1111/2041-210X.12970

Jactel, H., Koricheva, J., & Castagneyrol, B. (2019). Responses of forest insect pests to climate change: Not so simple. Current Opinion in Insect Science, 35, 103–108. https://doi.org/10.1016/j.cois.2019.07.010

Kindlmann, P., Dixon, A. F. G., & Dostálková, I. (2001). Role of ageing and temperature in shaping reaction norms and fecundity functions in insects. Journal of Evolutionary Biology, 14(5), 835–840. https://doi.org/10.1046/j.1420-9101.2001.00323.x

Kleynhans, E., Mitchell, K. A., Conlong, D. E., & Terblanche, J. S. (2014). Evolved variation in cold tolerance among populations of Eldana saccharina (Lepidoptera: Pyralidae) in South Africa. Journal of Evolutionary Biology, 27(6), 1149–1159. https://doi.org/10.1111/jeb.12390

Klock, C. J., & Chown, S. L. (2003). Resistance to temperature extremes in sub-Antarctic weevils: Interspecific variation, population differentiation and acclimation. Biological Journal of the Linnean Society, 78(3), 401–414. https://doi.org/10.1046/j.1095-8312.2003.00154.x

Kretschmer, M., Coumou, D., Agel, L., Barlow, M., Tziperman, E., & Cohen, J. (2018). More-persistent weak stratospheric polar vortex states linked to cold extremes. Bulletin of the American Meteorological Society, 99(1), 49–60. https://doi.org/10.1175/BAMS-D-16-0259.1

Lancaster, L. T. (2016). Widespread range expansions shape latitudinal variation in insect thermal limits. Nature Climate Change, 6(6), 618–621. https://doi.org/10.1038/nclimate2945

Lawrence, D. M., & Slater, A. G. (2010). The contribution of snow condition trends to future ground climate. Climate Dynamics, 34(7–8), 969–981. https://doi.org/10.1007/s00382-009-0537-4

Liebhold, A. M. (1995). Suitability of North American Tree Species to the Gypsy Moth: A Summary of Field and Laboratory Tests. U.S. Department of Agriculture, Forest Service, Northeastern Forest Experiment Station.

MacLean, H. J., Sørensen, J. G., Kristensen, T. N., Loeschcke, V., Beedholm, K., Kellermann, V., & Overgaard, J. (2019). Evolution and plasticity of thermal performance: An analysis of variation in thermal tolerance and fitness in 22 Drosophila species. Philosophical Transactions of the Royal Society B: Biological Sciences, 374(1778), 20180548. https://doi.org/10.1098/rstb.2018.0548

MacMillan, H. A., & Sinclair, B. J. (2011). Mechanisms underlying insect chill-coma. Journal of Insect Physiology, 57(1), 12–20. https://doi.org/10.1016/j.jinsphys.2010.10.004

MacQuarrie, C. J. K., *, Cooke, B. J., & Saint-Amant, R. (2019). The predicted effect of the polar vortex of 2019 on winter survival of emerald ash borer and mountain pine beetle. Canadian Journal of Forest Research, 49(9), 1165–1172. https://doi.org/10.1139/cjfr-2019-0115

Marshall, K. E., Gotthard, K., & Williams, C. M. (2020). Evolutionary impacts of winter climate change on insects. Current Opinion in Insect Science, 41, 54–62. https://doi.org/10.1016/j.cois.2020.06.003

Morgan, T. J., & Mackay, T. F. C. (2006). Quantitative trait loci for thermotolerance phenotypes in Drosophila melanogaster. Heredity, 96(3), 232–242. https://doi.org/10.1038/sj.hdy.6800786

PRISM Climate Group, Oregon State University, 30-Year Normals 1981-2010. https://prism.oregonstate.edu, data created 11 July 2012, accessed 7 Nov 2018.

Pureswaran, D. S., Roques, A., & Battisti, A. (2018). Forest insects and climate change. Current Forestry Reports, 4(2), 35–50. https://doi.org/10.1007/s40725-018-0075-6

Roland, J., & Matter, S. F. (2016). Pivotal effect of early-winter temperatures and snowfall on population growth of alpine Parnassius smintheus butterflies. Ecological Monographs, 86(4), 412–428. https://doi.org/10.1002/ecm.1225

Rossiter, M. C., Cox-Foster, D. L., & Briggs, M. A. (1993). Initiation of maternal effects in Lymantria dispar: Genetic and ecological components of egg provisioning. Journal of Evolutionary Biology, 6(4), 577–589. https://doi.org/10.1046/j.1420-9101.1993.6040577.x

Sharov, A. A., Leonard, D., Liebhold, A. M., Roberts, E. A., & Dickerson, W. (2002). “Slow the Spread”: A national program to contain the gypsy moth. Journal of Forestry, 100(5), 30–36. https://doi.org/10.1093/jof/100.5.30

Sinclair, B. J., Coello Alvarado, L. E., & Ferguson, L. V. (2015). An invitation to measure insect cold tolerance: Methods, approaches, and workflow. Journal of Thermal Biology, 53, 180–197. https://doi.org/10.1016/j.jtherbio.2015.11.003

Sinclair, B. J., & Roberts, S. P. (2005). Acclimation, shock and hardening in the cold. Journal of Thermal Biology, 30(8), 557–562. https://doi.org/10.1016/j.jtherbio.2005.07.002

Sinclair, B. J., Williams, C. M., & Terblanche, J. S. (2012). Variation in thermal performance among insect populations. Physiological and Biochemical Zoology, 85(6), 594–606. https://doi.org/10.1086/665388

Sisodia, S., & Singh, B. N. (2010). Influence of developmental temperature on cold shock and chill coma recovery in Drosophila ananassae: Acclimation and latitudinal variations among Indian populations. Journal of Thermal Biology, 35(3), 117–124. https://doi.org/10.1016/j.jtherbio.2010.01.001

Streifel, M. A., Tobin, P. C., Kees, A. M., & Aukema, B. H. (2019). Range expansion of Lymantria dispar dispar (L.) (Lepidoptera: Erebidae) along its north-western margin in North America despite low predicted climatic suitability. Journal of Biogeography, 46(1), 58–69. https://doi.org/10.1111/jbi.13474

Thompson, L. M., Faske, T. M., Banahene, N., Grim, D., Agosta, S. J., Parry, D., Tobin, P. C., Johnson, D. M., & Grayson, K. L. (2017). Variation in growth and developmental responses to supraoptimal temperatures near latitudinal range limits of gypsy moth Lymantria dispar (L.), an expanding invasive species. Physiological Entomology, 42(2), 181–190. https://doi.org/10.1111/phen.12190

Thompson, L. M., Powers, S. D., Appolon, A., Hafker, P., Milner, L., Parry, D., Agosta, S. J., & Grayson, K. L. (2021). Climate-related geographical variation in performance traits across the invasion front of a widespread non-native insect. Journal of Biogeography, 48(2), 405–414. https://doi.org/10.1111/jbi.14005

Thornton, M. M., Shrestha, R., Wei, Y., Thornton, P. E., Kao, S.-C., & Wilson, B. E. (2020). Daymet: Daily surface weather data on a 1-km grid for North America, Version 4. ORNL DAAC. https://doi.org/10.3334/ORNLDAAC/1840

Vavrus, S., Walsh, J. E., Chapman, W. L., & Portis, D. (2006). The behavior of extreme cold air outbreaks under greenhouse warming. International Journal of Climatology, 26(9), 1133–1147. https://doi.org/10.1002/joc.1301

Williams, C. M., Henry, H. A. L., & Sinclair, B. J. (2015). Cold truths: How winter drives responses of terrestrial organisms to climate change. Biological Reviews, 90(1), 214–235. https://doi.org/10.1111/brv.12105

Wu, Y., Molongoski, J. J., Winograd, D. F., Bogdanowicz, S. M., Louyakis, A. S., Lance, D. R., Mastro, V. C., & Harrison, R. G. (2015). Genetic structure, admixture and invasion success in a Holarctic defoliator, the gypsy moth (Lymantria dispar, Lepidoptera: Erebidae). Molecular Ecology, 24(6), 1275–1291. https://doi.org/10.1111/mec.13103

Yang, L. H., Postema, E. G., Hayes, T. E., Lippey, M. K., & MacArthur-Waltz, D. J. (2021). The complexity of global change and its effects on insects. Current Opinion in Insect Science, 47, 90–102. https://doi.org/10.1016/j.cois.2021.05.001

Zachariassen, K. E. (1985). Physiology of cold tolerance in insects. Physiological Reviews, 65(4), 799–832. https://doi.org/10.1152/physrev.1985.65.4.799

